# Classification of patients with COVID-19 by blood RNA endotype: A prospective cohort study

**DOI:** 10.1101/2023.06.22.546100

**Authors:** Jumpei Yoshimura, Yuki Togami, Takeshi Ebihara, Hisatake Mastumoto, Yumi Mitsuyama, Fuminori Sugihara, Haruhiko Hirata, Daisuke Okuzaki, Hiroshi Ogura

**Affiliations:** Department of Traumatology and Acute Critical Medicine, Osaka University Graduate School of Medicine, Suita, Japan; Center for Infectious Disease Education and Research, Osaka University; Division of Trauma and Surgical Critical Care, Osaka General Medical Center, Osaka, Japan; Core Instrumentation Facility, Immunology Frontier Research Center and Research Institute for Microbial Diseases, Osaka University, Osaka, Japan; Department of Respiratory Medicine and Clinical Immunology, Osaka University Graduate School of Medicine, Osaka, Japan; Laboratory of Human Immunology (Single Cell Genomics), WPI Immunology Research Center, Osaka University, Osaka, Japan; Genome Information Research Center, Research Institute for Microbial Diseases, Osaka University, Osaka, Japan

**Keywords:** SARS-CoV-2, phenotype, subtype, clustering, prognostic biomarker, COVID-19

## Abstract

**Background:** Although the development of vaccines has considerably reduced the severity of COVID-19, its incidence is still high. Hence, a targeted approach based on RNA endotypes of a population should be developed to help design biomarker-based therapies for COVID-19.

**Objectives:** We evaluated the major RNAs transcribed in blood cells during COVID-19 using PCR to further elucidate its pathogenesis and determine predictive phenotypes in COVID-19 patients.

**Study design:** In a discovery cohort of 40 patients with COVID-19, 26,354 RNAs were measured on day 1 and day 7. Five RNAs associated with disease severity and prognosis were derived. In a validation cohort of 153 patients with COVID-19 treated in the intensive care unit, we focused on prolactin (PRL), and toll-like receptor 3 (TLR3) among RNAs, which have a strong association with prognosis, and evaluated the accuracy for predicting survival of PRL-to-TL3 ratios (PRL/TLR3) with the areas under the ROC curves (AUC). The validation cohort was divided into two groups based on the cut-off value in the ROC curve with the maximum AUC. The two groups were defined by high PRL/TLR3 (n=47) and low PRL/TLR3 groups (n=106) and the clinical outcomes were compared.

**Results:** In the validation cohort, the AUC for PRL/TLR3 was 0.79, showing superior prognostic ability compared to severity scores such as APACHE II and SOFA. The high PRL/TLR3 group had a significantly higher 28-day mortality than the low PRL/TLR3 group (17.0% vs 0.9%, P<0.01).

**Conclusions:** A new RNA endotype classified using high PRL/TLR3 was associated with mortality in COVID-19 patients.

## Background

The global COVID-19 pandemic at the end of 2019 has claimed millions of lives and the toll continues to rise. Its unprecedented impact has caused extreme pressure on health systems and economies as well as a massive health crisis. The complexity of this crisis is compounded by “long COVID,” a condition in which symptoms persist after recovery, reducing the quality of life of survivors and placing an additional strain on healthcare resources [1].

Toll-like receptors (TLRs), a class of pattern recognition receptors, play an important role in determining the severity of COVID-19. SARS-CoV-2, classified as a pathogen-associated molecular patterns (PAMPs), interacts with TLR. This interaction initiates an intracellular signaling cascade that leads to the release of inflammatory cytokines such as IL-1, IL-6 and TNF-α [2]. An excessive immune response leads to a cytokine storm, which in turn triggers acute respiratory distress syndrome (ARDS) [1].

Recently, it has been reported that comprehensive analysis of biomolecular information using blood and other tissues can classify various diseases into detailed molecular forms called endotypes [3–5]. The real-time reverse transcription-polymerase chain reaction (RT-PCR) tests, which detect viral RNA, are mainly used for the diagnosis of COVID-19. This test detects COVID-19 viral RNA, making it possible to identify COVID-19 associated ARDS among diverse molecular patterns of ARDS [6]. However, it is insufficient to predict disease severity and provide insight into the host immune response to guide targeted immunological therapy.

The purpose of this study is to evaluate the blood genome endotypes of COVID-19 to understand the molecular pathogenesis of COVID-19.

## Objectives

In the present study, we aimed to investigate the transcribed RNA endotype signature in the blood cells of COVID-19 patients using a discovery and validation cohort. RNA sequencing (RNA-seq), reverse-transcription quantitative PCR (RT-qPCR), and bioinformatics analysis were applied to identify and examine the efficacy of the RNA endotypes in predicting the mortality and molecular pathogenesis of COVID-19.

## Study design

### Cohort data

In the present study, we used data from a discovery cohort (40 COVID-19 patients and 16 healthy volunteers) and a validation cohort (153 COVID-19 patients). The details of the cohorts are included in Supplemental Methods. An overview of patient flow, samples, and data analysis is presented in Fig. 1.

**Figure 1.**
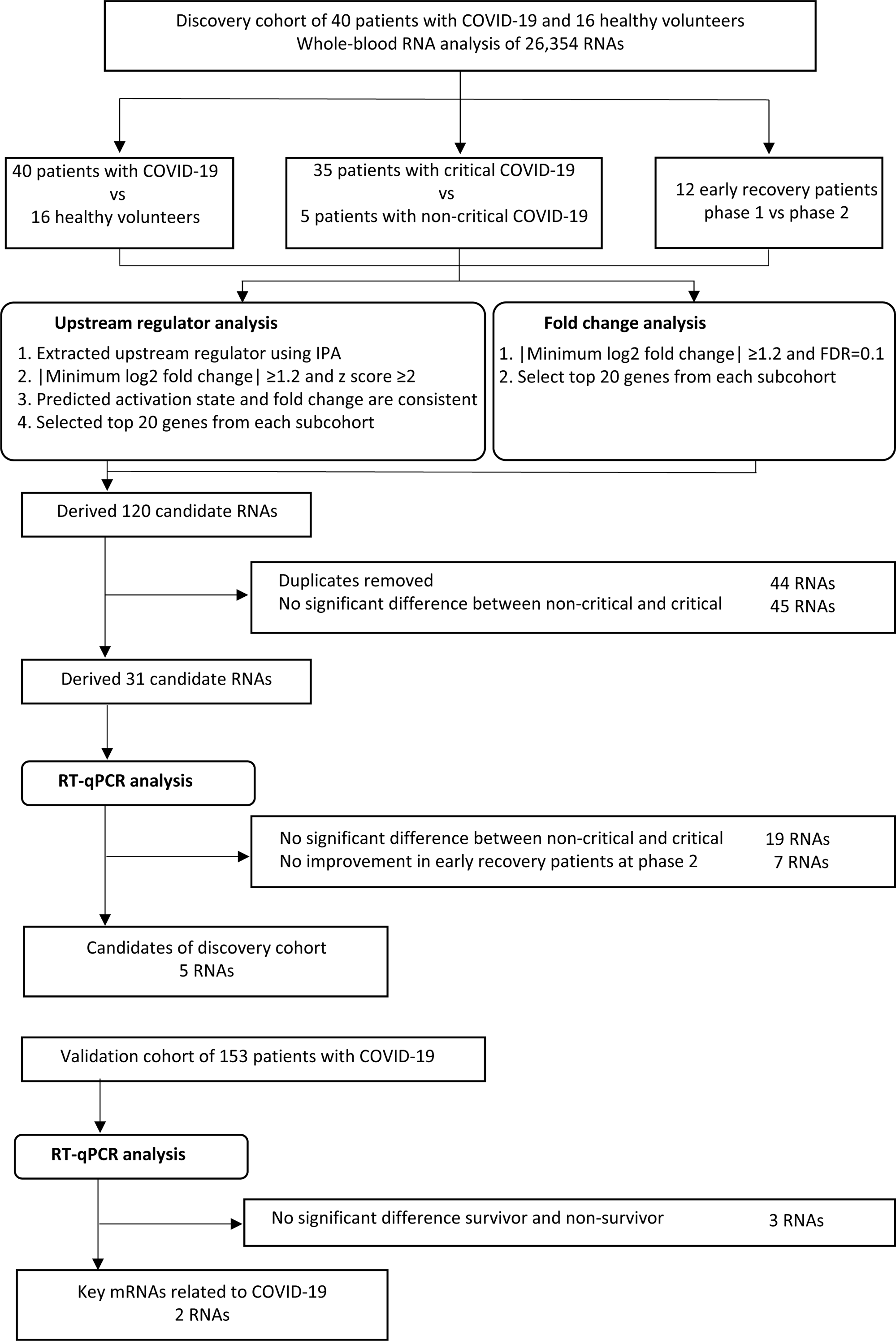
Patient cohorts, samples, and data analysis. COVID-19: Coronavirus disease 2019.

### Disease severity: Critical vs. non-critical

Acuity scores were based on the World Health Organization ordinal outcome scale [7]: A1, dead; A2, intubated, survived; A3, hospitalized with oxygen; A4, hospitalized without oxygen; A5, discharged. Disease severity was classified according to the maximum acuity score (acuity max 1 or 2). In this study, we defined ‘critical’ as acuity max 1 and 2 patients and ‘non-critical’ as acuity max 3, 4, and 5 patients.

### Sample collection timing: Phase 1 vs. phase 2

The timing of sample collection varied among the two cohorts. Therefore, we categorized them into the following two phases: phase 1 as days 1–2, and phase 2 as days 6–14. Day 1 referred to the day of admission to the hospital.

### Clinical outcome: Early recovery vs. late recovery

We defined the clinical outcome of patients treated with intermittent mandatory ventilation for ≤12 days or not as early recovery and that for intermittent mandatory ventilation for >12 days or 28-day non-survivors as late recovery, as per our previous study [8]. We divided the COVID-19 patients of the discovery cohort into two groups based on early and late recovery (Fig. 1).

### RNA sequencing (RNA-seq) and reverse-transcription quantitative PCR (RT-qPCR)

RNA-seq was performed as previously described [9]. Details are presented in Supplemental Methods. RT-qPCR was performed using a Biomark HD (Fluidigm) microfluidics system as previously described [10, 11], with some modifications. Details are presented in Supplemental Methods.

### Statistical analysis

The study size was derived as per a similar previous study [12]. A subset of different conditions was created to extract molecules expressed in clinically important phenotypes. In the discovery cohort, we created three subcohorts that could be related to disease progression: healthy volunteers and COVID-19 patients, critical and non-critical, and early recovery phase 1 and phase 2. In each subcohort, the differences in the expression of 26,354 RNAs were evaluated. Statistical analysis of RNA-seq data was performed as previously described [9]. Multidimensional scaling using the cmdscale command in R version 4.1.2 [13] was performed using log2-normalized RNA fragments per kilobase of transcript per million mapped read (FPKM) values to compare expression between healthy volunteers and COVID-19 patients. Volcano plot analyses were performed to identify significant changes in RNA expression between patients and healthy volunteers, critical and non-critical subcohorts, and phase 1 and phase 2 subcohorts of the early recovery patients. Significance was defined as |minimum log2 fold change| >1.2 and false discovery rate <0.1. The top 20 significant RNAs were used for further analysis.

To evaluate the upstream regulators of RNA expression, the data were subjected to upstream regulator analysis, and z-scores and P values were calculated using ingenuity pathway analysis (IPA; QIAGEN Inc., https://www.qiagen.com/us/products/discovery-and-translational-research/next-generation-sequencing/informatics-and-data/interpretation-content-databases/ingenuity-pathway-analysis/) [14]. The z-score predicts the activation state of the upstream regulator using RNA expression patterns of the downstream state of that regulator. The upstream regulator was considered activated at |z-score| >2 with P <0.05. The top 20 significant upstream regulators were used for further analysis.

Based on the results of statistical analysis of RNA-seq results, RT-qPCR of 31 RNAs was performed. The Wilcoxon rank sum test was used for analysis, and P <0.05 was considered to indicate statistical significance.

The 5 candidate RNAs from the discovery cohort were validated in the validation cohort. To identify molecules associated with the most important prognostic phenotype, the patients were divided into two groups, 28-day survivors and 28-day non-survivors. The 5 candidate RNAs of the discovery cohort were compared by the Wilcoxon rank sum test between the two groups. The RNAs that were significantly changed (P <0.05) between the survivors and non-survivors were extracted as key mRNAs. Receiver operator characteristic (ROC) curves were created using key mRNAs and PRL-to-TL3 ratios (PRL/TLR3). The areas under the ROC curves (AUCs) were calculated to evaluate the accuracy for predicting survival.

The validation cohort was divided into two groups based on the cut-off value in the ROC curve with the maximum AUC. The two groups were defined by their clinical phenotypes, namely high PRL/TLR3 (n = 47) and low PRL/TLR3 groups (n = 106) and the clinical data of the two phenotypes were compared. Fisher’s exact test and the Wilcoxon rank sum test were used to compare phenotypes based on baseline characteristics and 28-day outcomes. Missing values were not imputed in any analyses. Cumulative mortality was determined using Kaplan-Meier curves, and the phenotypes were compared using the log rank test. A two-sided P value <0.05 was considered to indicate statistical significance. For all statistical analyses, a fully scripted data management pathway was created within the R environment for statistical computing, version 4.0.2 (R Foundation for Statistical Computing, Vienna, Austria). Categorical variables are reported as numbers and percentages, and significance was calculated using χ2 or Fisher’s exact test. Continuous variables are described using mean and standard error or compared using the Mann-Whitney U test or Wilcoxon rank sum test and described using median and interquartile range (IQR) values.

## Results

### Candidate RNAs as prognostic biomarkers for COVID-19 from the discovery cohort

In the present study, we enrolled 40 COVID-19 patients and 16 healthy volunteers in the discovery cohort. The cohort comprised 35 critical and 5 non-critical patients; 12 critical patients were classified as early recovery patients and 23 as late recovery patients. Detailed characteristics of the patients and healthy volunteers are shown in Table 1. We selected 120 candidate RNAs based on fold change and upstream analyses of the three subcohorts (COVID-19 patients vs. healthy volunteers, critical patients vs. non-critical patients, and early recovery phase 1 patients vs. phase 2 patients) (Fig. S1, Table S1). Thirty-one RNAs exhibited significant change in expression between critical and non-critical patients (Fig. S2). The expression of these 31 RNAs was re-evaluated using RT-qPCR, which revealed significant change in the expression of five RNAs, namely interleukin-18R1 (IL18R1), galectin-2 (LGALS2), mitogen-activated protein kinase 14 (MAPK14), prolactin (PRL), and toll-like receptor 3 (TLR3), in the three subcohorts (Fig. S3). Thus, these five RNAs were investigated in the validation cohort (Fig. 1).

**Table 1.**
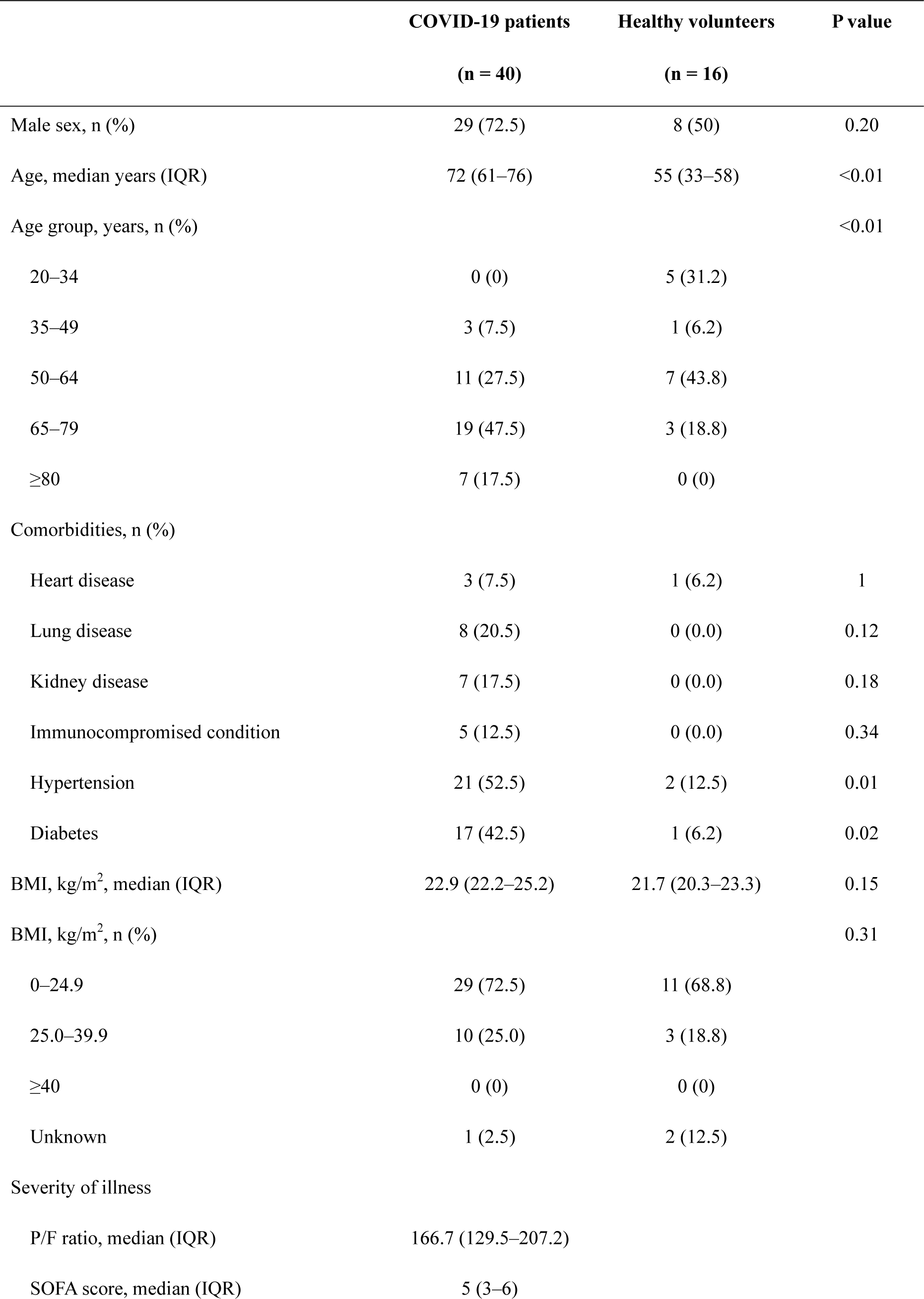
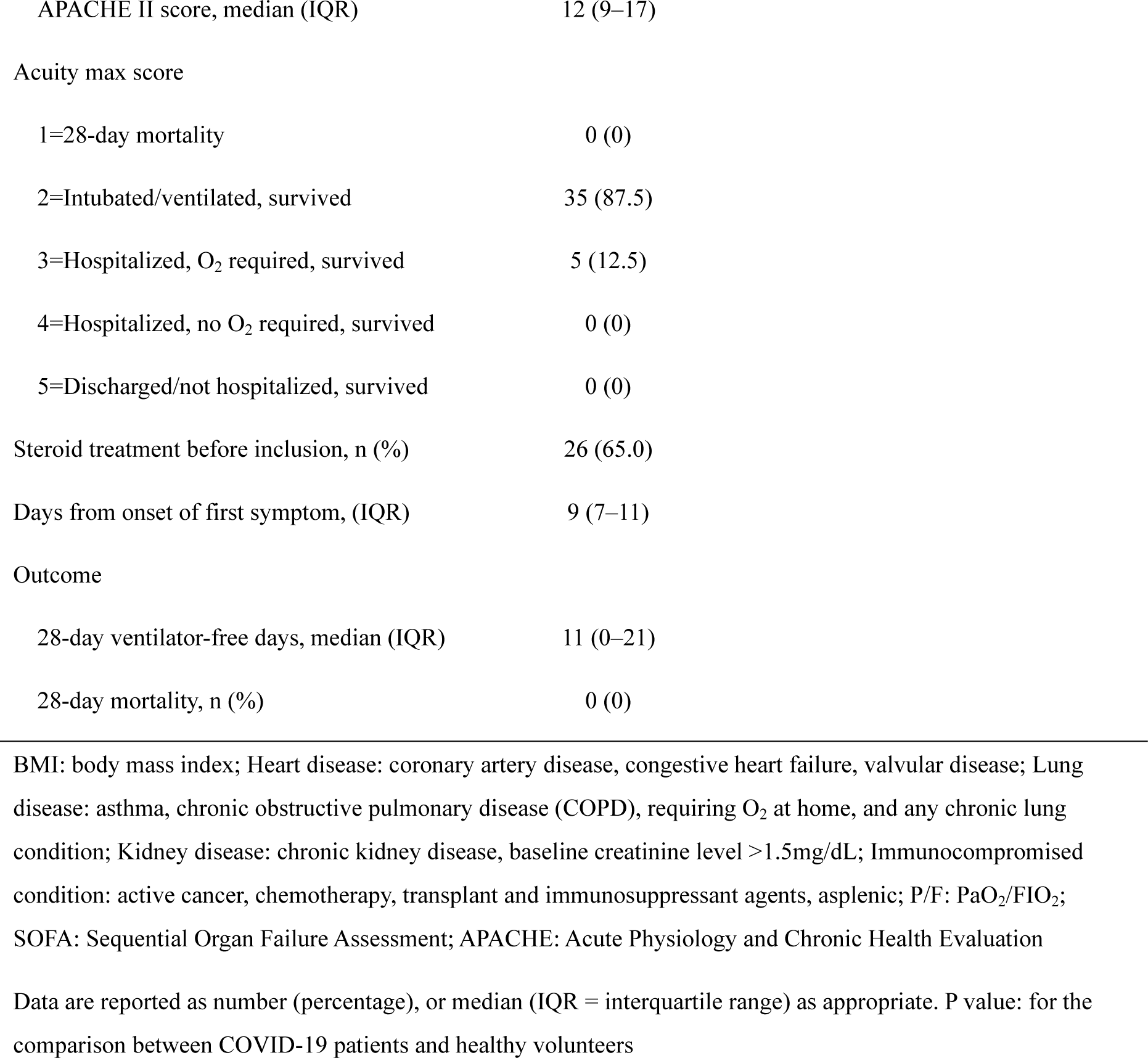
Clinical and demographic characteristics of COVID-19 patients and healthy volunteers in the discovery cohort

|  | COVID-19 patients<br>(n = 40) | Healthy volunteers<br>(n = 16) | P value |
| --- | --- | --- | --- |
| Male sex, n (%) | 29 (72.5) | 8 (50) | 0.20 |
| Age, median years (IQR) | 72 (61–76) | 55 (33–58) | <0.01 |
| Age group, years, n (%) |  |  | <0.01 |
| 20–34 | 0 (0) | 5 (31.2) |  |
| 35–49 | 3 (7.5) | 1 (6.2) |  |
| 50–64 | 11 (27.5) | 7 (43.8) |  |
| 65–79 | 19 (47.5) | 3 (18.8) |  |
| ≥80 | 7 (17.5) | 0 (0) |  |
| Comorbidities, n (%) |  |  |  |
| Heart disease | 3 (7.5) | 1 (6.2) | 1 |
| Lung disease | 8 (20.5) | 0 (0.0) | 0.12 |
| Kidney disease | 7 (17.5) | 0 (0.0) | 0.18 |
| Immunocompromised condition | 5 (12.5) | 0 (0.0) | 0.34 |
| Hypertension | 21 (52.5) | 2 (12.5) | 0.01 |
| Diabetes | 17 (42.5) | 1 (6.2) | 0.02 |
| BMI, kg/m<sup>2</sup>, median (IQR) | 22.9 (22.2–25.2) | 21.7 (20.3–23.3) | 0.15 |
| BMI, kg/m<sup>2</sup>, n (%) |  |  | 0.31 |
| 0–24.9 | 29 (72.5) | 11 (68.8) |  |
| 25.0–39.9 | 10 (25.0) | 3 (18.8) |  |
| ≥40 | 0 (0) | 0 (0) |  |
| Unknown | 1 (2.5) | 2 (12.5) |  |
| Severity of illness |  |  |  |
| P/F ratio, median (IQR) | 166.7 (129.5–207.2) |  |  |
| SOFA score, median (IQR) | 5 (3–6) |  |  |

|  |  |
| --- | --- |
| APACHE II score, median (IQR) | 12 (9–17) |
| Acuity max score |  |
| 1=28-day mortality | 0 (0) |
| 2=Intubated/ventilated, survived | 35 (87.5) |
| 3=Hospitalized, O<sub>2</sub> required, survived | 5 (12.5) |
| 4=Hospitalized, no O<sub>2</sub> required, survived | 0 (0) |
| 5=Discharged/not hospitalized, survived | 0 (0) |
| Steroid treatment before inclusion, n (%) | 26 (65.0) |
| Days from onset of first symptom, (IQR) | 9 (7–11) |
| Outcome |  |
| 28-day ventilator-free days, median (IQR) | 11 (0–21) |
| 28-day mortality, n (%) | 0 (0) |

BMI: body mass index; Heart disease: coronary artery disease, congestive heart failure, valvular disease; Lung disease: asthma, chronic obstructive pulmonary disease (COPD), requiring O<sub>2</sub> at home, and any chronic lung condition; Kidney disease: chronic kidney disease, baseline creatinine level >1.5mg/dL; Immunocompromised condition: active cancer, chemotherapy, transplant and immunosuppressant agents, asplenic; P/F: PaO<sub>2</sub>/FIO<sub>2</sub>; SOFA: Sequential Organ Failure Assessment; APACHE: Acute Physiology and Chronic Health Evaluation
Data are reported as number (percentage), or median (IQR = interquartile range) as appropriate. P value: for the comparison between COVID-19 patients and healthy volunteers

### Validation of five candidate RNAs using RT-qPCR

To ascertain the robustness of the five RNAs as candidate biomarkers for COVID-19 prognosis, we assessed their expression in an independent validation cohort using RT-qPCR. We included 153 COVID-19 patients in the validation cohort, which comprised 145 critical and 8 non-critical patients; 9 patients (5.9%) died within 28 days after ICU admission. The characteristics of the patients are presented in Table S2. RT-qPCR analysis revealed that the RNA expression of PRL significantly increased and that of TLR3 significantly decreased in 28-day non-survivors compared with that in 28-day survivors (Fig. 2). Therefore, PRL and TLR3 were established as the two key RNAs related to the prognosis of COVID-19 patients.

**Figure 2.**
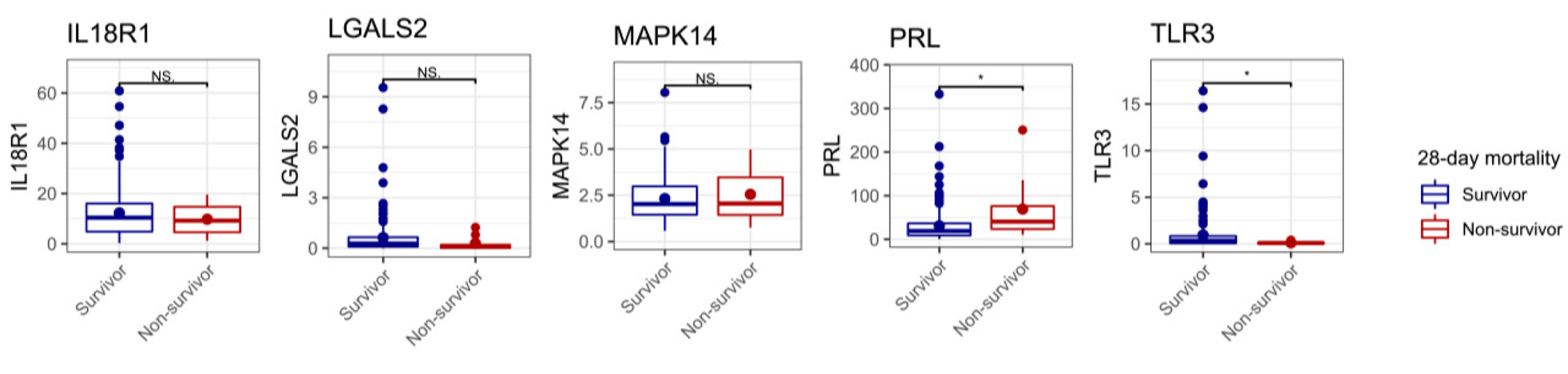
Identification of two RNAs related to mortality of COVID-19 patients from the validation cohort. Among the five candidate RNAs selected from the analysis of the discovery cohort, the expression of PRL was significantly increased and that of TLR3 was significantly decreased in the 28-day non-survivor subcohort compared with that in the 28-day survivor subcohort. NS, not significant. * P<0.05.

### RNA endotypes for discriminating COVID-19 patients with poor vs. good prognosis

Considering the characteristics of the expression changes of the two RNAs in patients with poor prognosis, we evaluated the ratio of PRL mRNA expression to that of TLR3 (PRL/TL3) as a biomarker of RNA phenotype. The performance of several protein markers (C-reactive protein, [CRP] and fibrin degradation product [FDP]) and severity scores (Acute Physiology and Chronic Health Evaluation II [APACHE II] and Sequential Organ Failure Assessment [SOFA]) were compared with that of PRL/TLR3 using ROC curve analysis. The AUC for PRL/TLR3 was 0.79 (95% confidence interval, 0.69–0.89), whereas those of CRP and FDP were 0.48 (0.29–0,66) and 0.66 (0.48–0.84), respectively. The AUCs for APACHE II and SOFA severity scores were 0.70 (0.58–0.81) and 0.49 (0.26–0.71), respectively. Therefore, PRL/TLR3 was more suitable for the discrimination of prognosis than the protein biomarkers and severity scores (Fig. 3). Hence, based on the ROC curve analysis, the threshold for PRL/TLR3 was set at 294.8.

**Figure 3.**
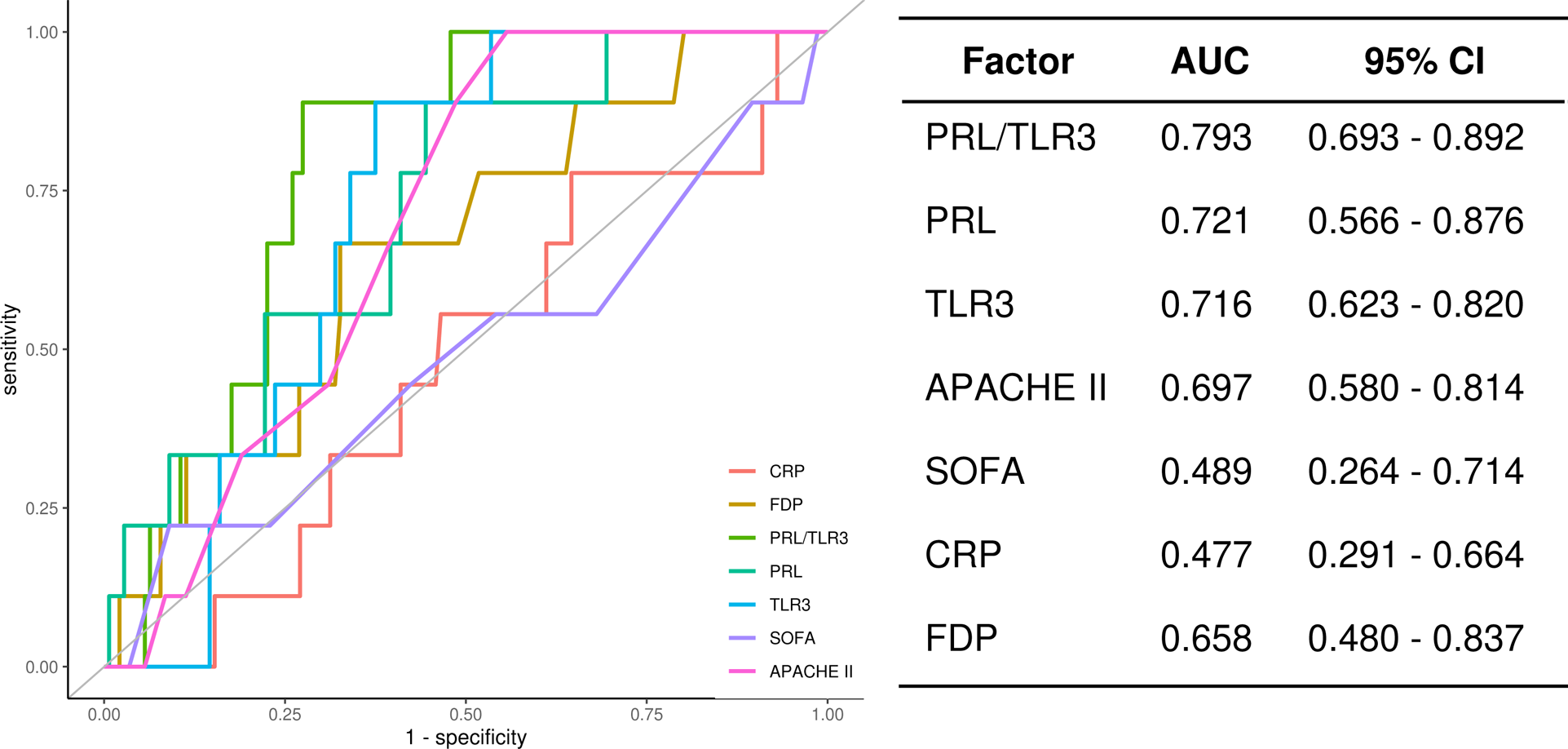
Prediction of mortality and clinical outcome. APACHE II, Acute Physiology and Chronic Health Evaluation II; AUC, area under the curve; CI, confidence interval; CRP, C-reactive protein; FDP, fibrin degradation products; PRL, prolactin; SOFA, Sequential Organ Failure Assessment; TLR3, Toll-like receptor 3.

### Comparison of patient outcomes based on PRL/TLR3

We divided 153 patients into two groups, namely high PRL/TLR3 (n = 47) and low PRL/TLR3 groups (n = 106) and compared patient outcomes by the groups. The RNA expression pattern of PRL and TLR3 in each group is shown in Fig. 4. The RNA endotypes did not exhibit an association with age, sex, comorbidities, PaO_2_/FIO_2_ (P/F ratio), or SOFA and APACHE II scores at the time of admission of the patients to the ICU (Table 2, Fig. 5a).

**Figure 4.**
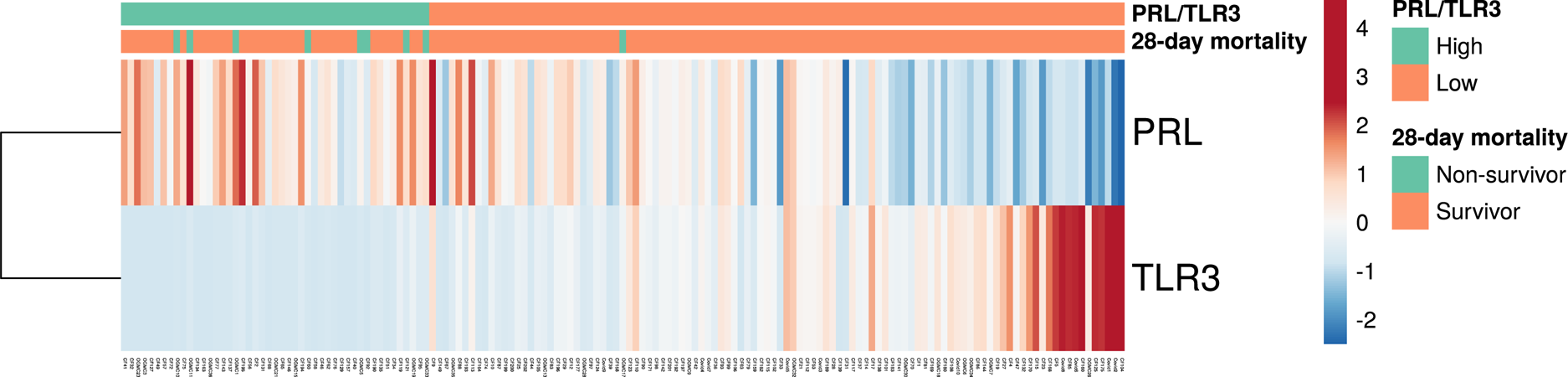
RNA expression pattern of PRL and TLR3. PRL, prolactin; TLR3, Toll-like receptor 3.

**Figure 5.**
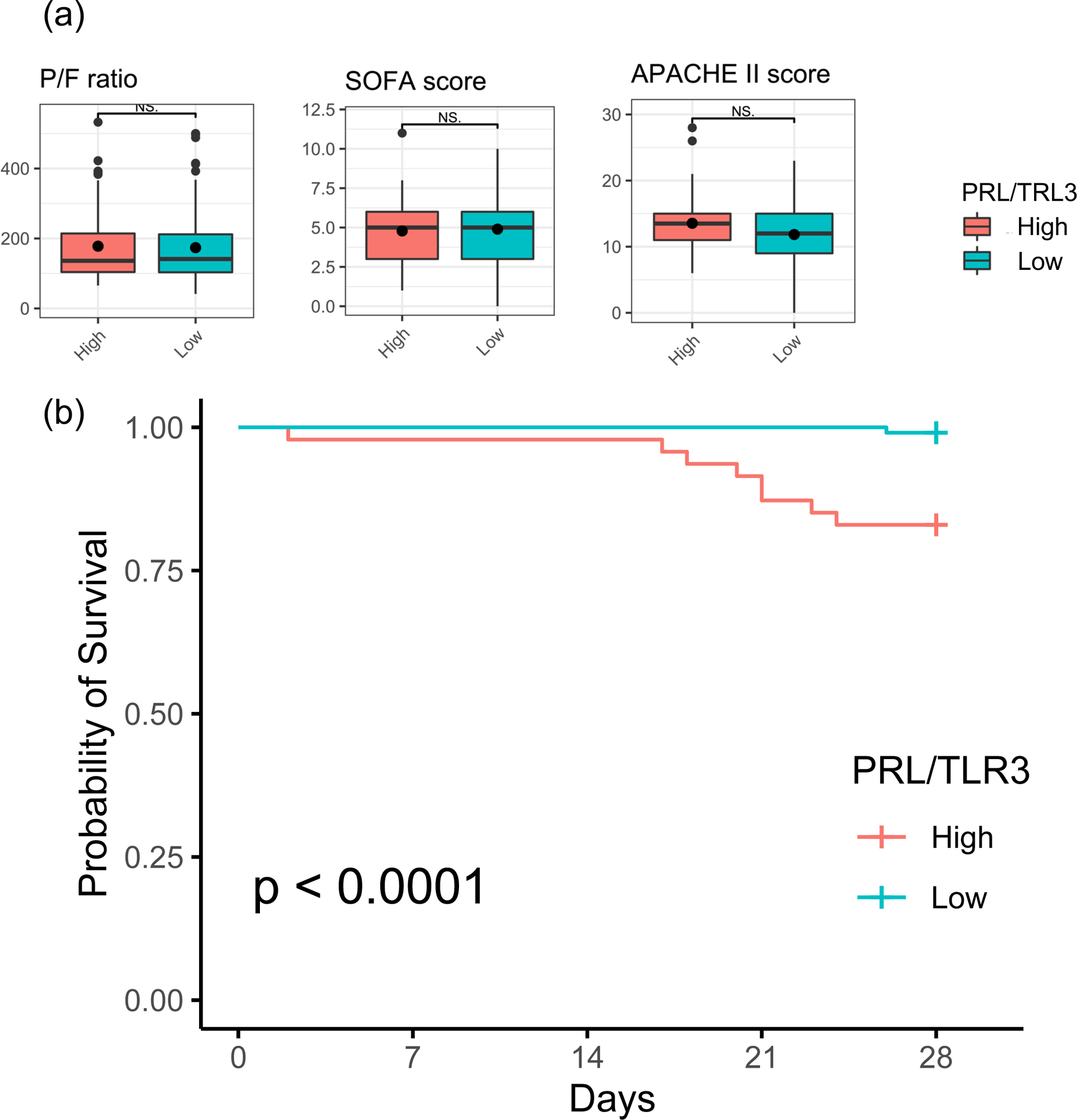
Survival analysis. (a) The RNA endotypes did not show an association with PaO_2_/FIO_2_ (ratio) or SOFA and APACHE II scores at admission to the ICU. (b) Kaplan-Meier analysis showed an association between the RNA endotype and 28-day mortality. APACHE II: Acute Physiology and Chronic Health Evaluation II; NS: not significant; SOFA: Sequential Organ Failure Assessment.

**Table 2.** Clinical and demographic characteristics of COVID-19 patients according to PRL/TLR3 ratio

|  | High PRL/TLR3<br>(n = 47) | Low PRL/TLR3<br>(n = 106) | P value |
| --- | --- | --- | --- |
| Male sex, n (%) | 38 (80.9) | 81 (76.4) | 0.69 |
| Age, years, median (IQR) | 60 (54–70) | 60 (52–67) | 0.31 |
| Age group, years, n (%) |  |  | 0.66 |
| 20–34 | 1 (2.1) | 6 (5.7) |  |
| 35–49 | 6 (12.8) | 15 (14.2) |  |
| 50–64 | 21 (44.7) | 53 (50.0) |  |
| 65–79 | 17 (36.2) | 27 (25.5) |  |
| ≥80 | 2 (4.3) | 5 (4.7) |  |
| Comorbidities, n (%) |  |  |  |
| Heart disease | 3 (6.4) | 8 (7.5) | 1 |
| Lung disease | 4 (8.5) | 7 (6.6) | 0.94 |
| Kidney disease | 4 (8.5) | 5 (4.7) | 0.58 |
| Immunocompromised condition | 3 (6.4) | 2 (1.9) | 0.34 |
| Hypertension | 15 (31.9) | 36 (34.0) | 0.95 |
| Diabetes | 12 (25.5) | 43 (40.6) | 0.11 |
| BMI, kg/m<sup>2</sup>, median (IQR) | 25.2 (22.6–28.3) | 26.5 (23.4–29.4) | 0.53 |
| BMI, kg/m<sup>2</sup>, n (%) |  |  | 0.53 |
| 0–24.9 | 20 (42.6) | 35 (33.0) |  |
| 25.0–39.9 | 23 (48.9) | 56 (52.8) |  |
| ≥40 | 0 (0.0) | 2 (1.9) |  |
| Unknown | 4 (8.5) | 13 (12.3) |  |
| Laboratory data at inclusion |  |  |  |
| White blood cells (10<sup>3</sup>/L) | 9600 (6800–12720) | 7970 (5480–10680) | 0.04 |
| Platelet count ($10^3/\text{L}$) | 19.3 (14.6–25.1) | 19.3 (15.6–255.8) | 0.47 |
| FDP ($\mu\text{g/mL}$) | 4.7 (3.7–12.4) | 4.7 (3.2–11.0) | 0.35 |
| Creatinine ($\text{mg/dL}$) | 0.7 (0.6–0.8) | 0.8 (0.6–1.0) | 0.39 |
| Bilirubin ($\text{mg/dL}$) | 0.6 (0.5–0.8) | 0.6 (0.4–0.8) | 0.77 |
| LDH ($\text{IU/L}$) | 520 (398–688) | 458 (373–646) | 0.30 |
| CRP ($\text{mg/dL}$) | 7.1 (3.3–6.0) | 5.0 (3.0–6.0) | 0.88 |
| Severity of illness |  |  |  |
| P/F ratio | 135.3 (102.9–211.2) | 142.6 (103.6–213.4) | 0.84 |
| SOFA score, median (IQR) | 5 (3–6) | 5 (3–6) | 0.80 |
| APACHE II score, median (IQR) | 14 (11–15) | 12 (9–15) | 0.80 |

BMI; body mass index; CRP: c-reactive protein; Heart disease: coronary artery disease, congestive heart failure, valvular disease; Lung disease: asthma, chronic obstructive pulmonary disease (COPD), requiring $\text{O}_2$ at home, and any chronic lung condition; Kidney disease: chronic kidney disease, baseline creatinine $>1.5$ $\text{mg/dL}$ ; Immunocompromised condition: active cancer, chemotherapy, transplant and immunosuppressant agents, asplenic; LDH: lactate dehydrogenase; P/F: $\text{PaO}_2/\text{FIO}_2$ ; SOFA: Sequential Organ Failure Assessment; APACHE: Acute Physiology and Chronic Health Evaluation
Data are reported as number (percentage) or median (IQR, interquartile range) as appropriate. P value: comparison between high and low group

The cumulative incidence of mortality at 28 days was 17.0% (n = 8) and 0.9% (n = 1) in the PRL/TLR3 high and low groups, respectively. Kaplan-Meier analysis revealed an association between RNA endotype and 28-day mortality (P<0.01) (Fig. 5b). Hospital mortality was significantly higher in the PRL/TLR3 high group than in the PRL/TLR3 low group (23.4% vs. 9.4%, P=0.04) (Table 3). However, we did not observe a difference in 28-day ventilator-free days and the duration of ICU stay between the two groups (P = 0.54 and P = 0.99, respectively).

**Table 3.**
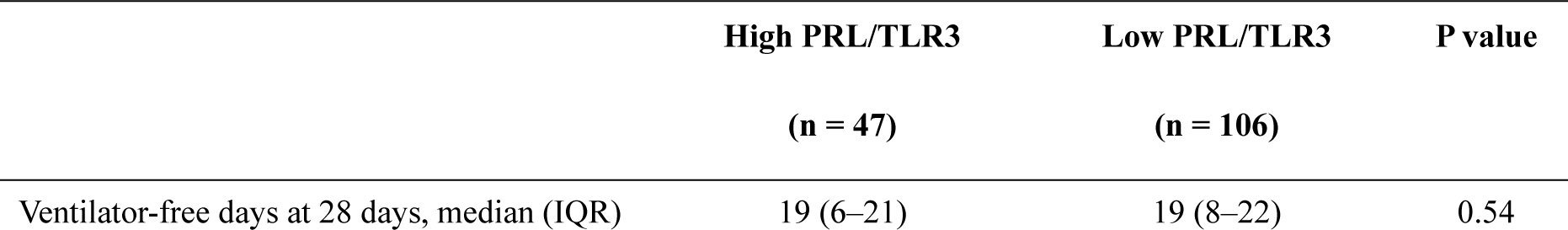
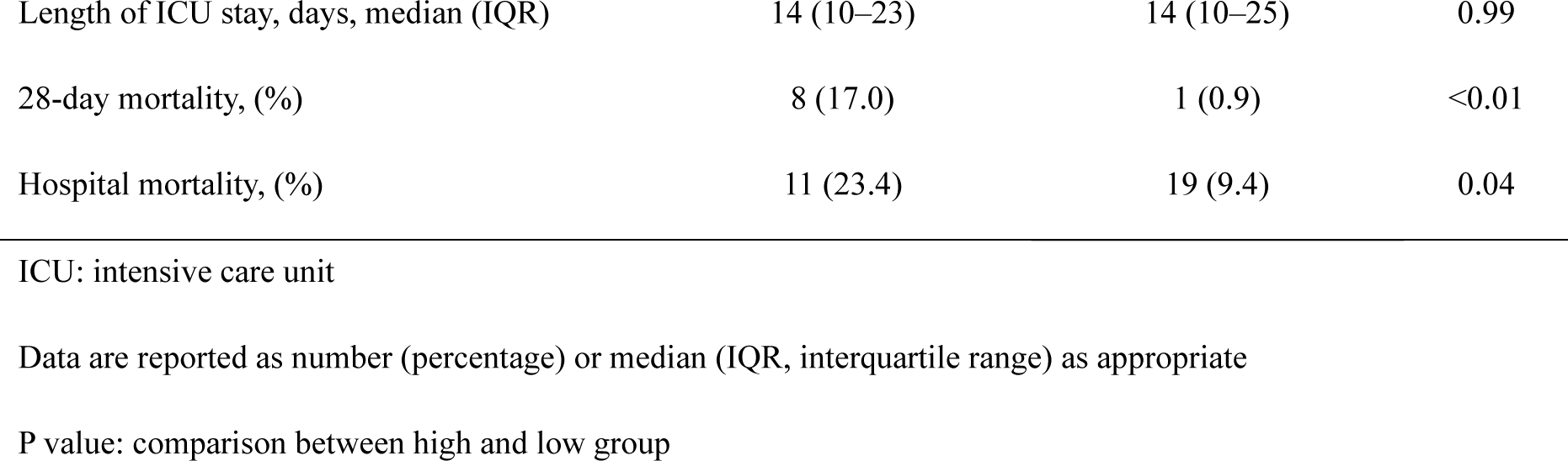
Clinical outcomes of COVID-19 patients according to PRL/TLR3 ratio

|  | High PRL/TLR3<br>(n = 47) | Low PRL/TLR3<br>(n = 106) | P value |
| --- | --- | --- | --- |
| Ventilator-free days at 28 days, median (IQR) | 19 (6–21) | 19 (8–22) | 0.54 |
| Length of ICU stay, days, median (IQR) | 14 (10–23) | 14 (10–25) | 0.99 |
| 28-day mortality, (%) | 8 (17.0) | 1 (0.9) | <0.01 |
| Hospital mortality, (%) | 11 (23.4) | 19 (9.4) | 0.04 |

ICU: intensive care unit
Data are reported as number (percentage) or median (IQR, interquartile range) as appropriate
P value: comparison between high and low group

## Discussion

In the present study, we identified high PRL/TLR3 and low PRL/TLR3 endotypes of COVID-19 patients in two heterogeneous cohorts using whole-blood samples of the patients at the time of admission to the ICU. We elucidated that the RNA endotypes of COVID-19 patients could accurately predict patient mortality; these endotypes were not easily discernible by clinical characteristics.

The present findings based on the RNA expression of PRL and TLR3 could have pathophysiological implications. PRL is a pituitary hormone involved in lactation, luteal functioning, and reproduction [15]. Moreover, leukocytes produce PRL, which plays a major role in immune responses [16–18]. PRL strongly affects the innate and adaptive immune responses, exerting an immunomodulatory effect at the early stages of T-cell activation and increasing the secretion of IFN-γ [17]. It also promotes cytokine production by monocytes and activates the STAT1 and MAPK pathways in granulocytes [19]. Some of these responses were also observed in the present study (Table S1, Fig. S2). Although various studies have reported elevated PRL expression and exacerbation of viral infections such as those caused by human immune deficiency virus and hepatitis C virus, the mechanisms underlying the high PRL expression in COVID-19 are poorly understood [20]. The results of our study suggest the potential role of PRL as a pro-inflammatory agent in COVID-19.

TLR3 plays a crucial role in the antiviral response against most viruses based on its ability to sense double-stranded RNA, a common intermediate of replication among many viruses [21]. A protective role of TLR3 has been reported in infections caused by organisms closely related with COVID-19 viruses, such as SARS-CoV-1 and Middle East respiratory syndrome coronavirus (MERS-CoV) [22]. Moreover, TLR3 mutants are predisposed to COVID-19 and influenza infection severity and associated mortality [23, 24]. Therefore, the association between reduced TLR3 expression and mortality revealed in the present study may reflect the consequences of impaired immune defense against COVID-19.

The identification of potential risk factors that predict the disease course will be useful for healthcare professionals to efficiently triage patients, personalize treatment, monitor clinical progress, and allocate proper resources at all levels of care to mitigate morbidity and mortality. Prognostic predictions of COVID-19 based on patient background and laboratory test results have been reported, but their accuracy is insufficient [25–27]. In the present study, the prognostic value of the RNA endotypes was higher than that of the existing clinical parameters, suggesting that the investigation of RNA expression in whole-blood samples may accurately assess pathophysiological changes that cannot be determined using clinical parameters. Furthermore, the differences in prognosis and therapeutic efficacy depending on RNA endotypes have been reported even in acute diseases, such as sepsis [3, 28]. Therefore, the accumulation of whole-blood RNA data of COVID-19 patients may reveal differences in response of critically ill patients to various therapies [29].

The study has a limitation. During the study period, SARS-CoV-2 clade B was mainly prevalent, with a bias toward it. Moreover, this clade was not identified in all cases. Therefore, this study does not elucidate the association between viral strain types and RNA endotypes. However, our study has several strengths over previous studies on RNA endotypes [3, 28]. We improved the accuracy of quantification by retesting the candidate genes obtained by RNA-seq using RT-qPCR. We also validated our results using multiple cohorts; therefore, the results should be considered robust.

Therefore, in the present study, we identified two key RNAs (PRL, TLR3) associated with the prognosis of COVID-19. A new RNA endotype classified using high PRL/TLR3 was associated with mortality in COVID-19 patients. This can potentially lead to the development of therapeutic interventions based on RNA endotypes in the future.

## Acknowledgments

We appreciate the cooperation of the patients, families, and healthy volunteers involved in this study. We also thank the medical staff for their cooperation. We acknowledge Shinichiro Kameoka and the NGS core facility of the Genome Information Research Center at the Research Institute for Microbial Diseases of Osaka University for their support in RNA sequencing and data analysis.

## Ethics approval

The study was performed in accordance with the Declaration of Helsinki and was approved by the institutional review board of Osaka University Hospital (Permit Number: 885 [Osaka University Critical Care Consortium Novel Omix Project; Occonomix Project]).

## Author contributions

JY conceived and designed this study, acquired and analyzed the data, and wrote the manuscript. HM helped with the study design and data interpretation and conducted the literature review. YT, TE, YM, and HH contributed to data acquisition. FS and DO contributed to data analysis. HO conducted the literature review.

## Data availability

The raw RNA sequence data of this study have been submitted under the Gene Expression Omnibus accession numbers GSE 182152 and GSE 179850.

## Informed consent and patient details

Informed consent was obtained from the patients or their relatives and the healthy volunteers for the collection of blood samples. Anonymized patient information was used, and no identifiable image was included in this study.

